# Cultivating Future Materials: Artificial Symbiosis for Bulk Production of Bacterial Cellulose Composites

**DOI:** 10.1101/2025.04.23.650277

**Authors:** Kui Yu, Sing Teng Chua, Alyssa Smith, Alison G. Smith, Tom Ellis, Silvia Vignolini

**Affiliations:** Yusuf Hamied Department of Chemistry, University of Cambridge, Cambridge CB2 1EW, United Kingdom; Department of Plant Sciences, University of Cambridge, Cambridge CB2 3EA, United Kingdom; Imperial College Centre for Synthetic Biology, London, SW7 2AZ, United Kingdom; Department of Bioengineering, Imperial College London, London, SW7 2AZ, United Kingdom; Sustainable and Bio-inspired Materials, Max Planck Institute of Colloids and Interfaces, Potsdam 14476, Germany

**Keywords:** bacterial cellulose, microalgae, living materials, co-culture, immobilization, biofilm, bacterial cellulose nanocrystals

## Abstract

Symbiotic relationships between micro-organisms are the basis of all global ecosystems. Here we extrapolate this concept for novel material fabrication by creating artificial symbiotic relationships between species that are usually not grown synergistically in nature. Specifically, we combine the cellulose-producing bacterium *Komagataeibacter hansenii* and the green microalga *Chlamydomonas reinhardtii* to obtain bulk growth of bacterial cellulose. Usually, bacterial cellulose is produced as floating pellicles at the air-liquid interface of the growing media, because free oxygen, together with the nutrients in the culture medium, is required for the bacteria to synthesise the cellulose fibres. In the co-culture, bacterial cellulose production can be achieved in bulk beyond the restriction of the liquid/air interface as the motile microalgae act as oxygen-generating sites within the culture medium. In exchange, the highly porous and mechanically robust scaffold provided by the cellulose allows the algal community to form a healthy biofilm, which can reach several centimetres thick. We demonstrate that our growing platform allows the simultaneous production of bulk bacterial cellulose in static incubation conditions, taking up an arbitrary and yet tunable 3D shape, dependent on the geometry of the culture vessel and with a relatively high biomass output.

## Introduction

The convergence of synthetic biology and materials science has ushered in a revolutionary way to a new class of materials called living materials, comprising living entities or communities of cells (such as bacteria^1^, microalgae^2,3^, yeast^4,5^ and fungi^6,7^) seamlessly integrated into self-regenerating matrices or artificial scaffolds.^8^ In this context, bacterial cellulose (BC) offers excellent opportunities for developing new engineered living materials with easy and cheap self-healing properties^9^. BC is produced at high yields by *Acetobacter*, and it presents a compelling pathway for the development of cost-effective materials for a wide range of applications (healthcare, biotechnology and electronic systems) in a local and circular economy to develop composites^4,9-15^ when compared to plant-extracted cellulose ^16-18^.

BC can be obtained by bacterial fermentation under mild temperatures (25-30 °C)^19^. This remarkable material owes its exceptional mechanical robustness^10^ to the high crystallinity of the cellulose polymer constituting the individual fibres and their layered arrangement^17^. Beyond its mechanical prowess, BC has garnered attention due to its sustainable production and functional bio-composite attributes. These include high purity, elevated porosity, excellent water retention, rapid production from living cells, sustainable production methods, self-regenerative capabilities, biocompatibility and biodegradability^20,21^.

However, while BC is potentially a much more sustainable source of nanocellulose when compared to plant-extracted ones from timber waste or cotton for example, it remains challenging to scale at comparable volumes. This is mainly because of high costs, as BC cannot be grown in traditional bioreactors, due to a lack of free oxygen^22^, since the cellulose pellicle forms only at the air-liquid interface, and during static incubation, oxygen diffusion from air into the depth of the culture medium is limited^13,14,23^. To overcome such limitations and develop a fully three-dimensional (3D) BC composite, several labour-intensive approaches, such as aerosol-based spraying^10,12^ and day-by-day shaking^23^ have been developed. The aerosol spraying method involves spraying a growth medium mixed with bacteria and functional components, such as graphene oxide (GO), as a gas mixture in a sealed bioreactor^10,12^. While such an approach increases the oxygen availability around the bacteria, aerosol spray requires very specialized equipment and poses challenges for large-scale production, such as potential contamination and inhomogeneity due to the sprayed gas. Providing simple agitation^23^ promotes BC production by improving oxygen diffusion into the medium and also leads to structural changes weakening the overall performance of the final composites, (such as mm-scale BC spheroids or stratification of the BC films^9,24^). To go beyond 2D pellicles, techniques, including foam templating^25^ and imprinting^26^, have also been developed to grow 3D BC-based composites^27^. However, the production of BC-based composites^14,27^ with adjustable 3D geometries by direct incubation^4,9^ remains challenging for truly scalable static systems.

Here, we propose a simple strategy to grow in bulk BC-based living composites with tuneable geometries by developing an artificial symbiosis between the cellulose-producing *K. hansenii*^19,28^ bacteria and the chlorophyte microalga *C. reinhardtii*^29,30^, as depicted in Figure 1a. Such symbiosis is enhanced by incorporating the growing medium with cellulose nanocrystals (CNCs)^31^. Such BC-derived colloidal particles enable the microalgal cells to remain dispersed homogeneously within the medium, acting as oxygen-generating sites to promote the production of the BC nanofiber network that surrounds them (Fig. 1b, c). As a result, simple incubation with little external intervention is necessary to grow materials with flexible geometries (Fig. 1d) in a sustainable and scalable approach. Simultaneously, the presence of the BC provides support to the healthy growth of a dense algal culture of several centimetres thick. We, therefore, believe that our methodology provides a sustainable and programmable way to develop mechanically robust BC-based bio-composites for multiple applications, including fibre-reinforced composites^32^, engineered living materials that can also be exploited in the contact of photobioreactors^32,3^, as it is possible to achieve both high cell viability (83%) and excellent mechanical robustness. Our symbiotic system also outperforms the properties of conventionally developed living materials such as artificial hydrogels^32,33^, where it remains challenging to overcome the trade-off between the problem of diffusion limitation (leading to low cell viability) and mechanical robustness with traditional crosslinking approaches^3,29,32,33^.

**Fig. 1.**
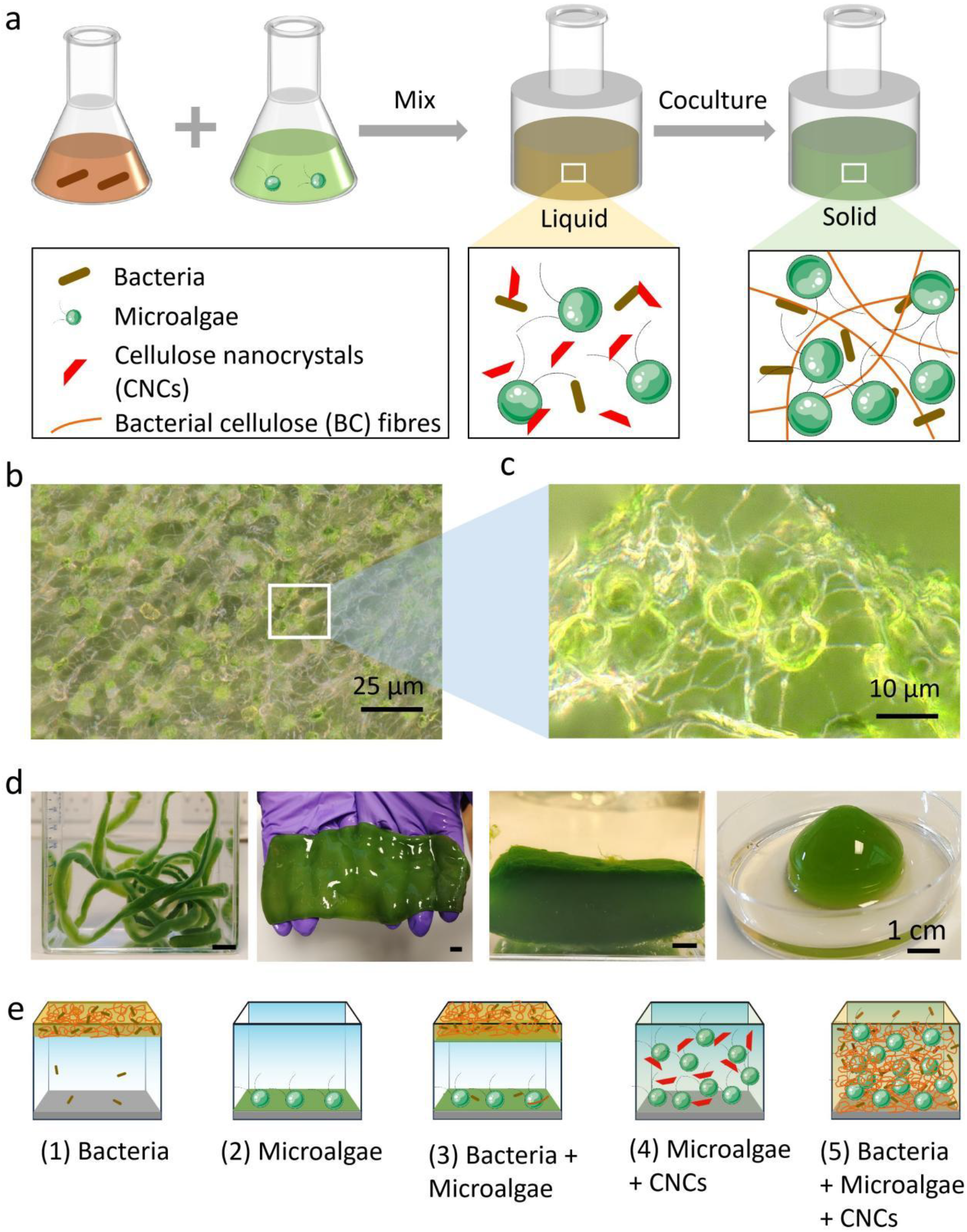
Generating BC-based bio-composites from coculture of bacteria (*K. hansenii*) and microalgae (*C. reinhardtii*). **a**, Schematic showing the fabrication procedure of BC-based bio-composites. **b**, Optical microscopic image for the cross-section of BC-based bio-composites, representing the successful incorporation of microalgal cells inside the BC layered structure. **c**, Optical microscopic image of microalgae morphology in the BC-based bio-composites, showing the cell-fibre interaction. **d**, BC-based bio-composites with tuneable geometries. **e**, Schematic for the status of bacteria culture, microalgae culture, bacteria and microalgae coculture, microalgae and CNCs mixture, and bacteria and microalgae coculture in the presence of CNCs under static incubation. The labels relating to bacteria, microalgae, CNCs and BC fibres in Fig. 1e are consistent with those of Fig. 1a.

## Results and Discussion

BC is generally produced by inoculating cellulose-producing bacteria, such as *K. hansenii* into a Hestrin-Schramm (HS) medium^19^. As the production of BC requires oxygen, the BC is formed as a solid flat pellicle at the air-liquid interface under static incubation (Fig. 1e(1), b, Supplementary Fig. 1a), due to the limited gaseous diffusion across the liquid medium^22^. While the incorporation of oxygen-generating microalgae serves as a potential solution, simply introducing oxygen-generating sites by adding microalgae to the liquid, as proposed in previous studies^14^, only allows for effective co-culture during agitation to increase oxygen diffusion and prevents sedimentation of the algae. In contrast, in static incubation, the microalgae cells sediment before the production of bacterial cellulose begins. We observed that during the algal sedimentation time (about 1 day in static incubation (Fig. 1e(2), Supplementary Fig. 1b), it is not possible to build large enough volumes of BC to encapsulate algal cells, which instead form a biofilm attached at the bottom of BC pellicle (Fig. 1e(3), c, Supplementary Fig. 2)^14^. Agitation of culture suspension, on the other hand, disrupted the formation of interconnected cellulosic network^34^.

To avoid microalgae sedimentation, we introduced CNCs in the culture medium. Specifically, we used BC-derived CNCs (B-CNCs) obtained by acid hydrolysis from freeze-dried BC fibres as previously reported in the literature^35^, although the same results can be obtained with most commonly used plant-derived CNCs (Supplementary Fig. 3). CNCs are rod-shaped, negatively charged nanoparticles that are colloidally stable in water suspensions. More details about the size of the used CNCs are provided in Supplementary Fig. 4. We observed that the addition of the CNCs to the microalgal culture prevented the sedimentation of cells (Fig. 1e(4)). We propose that such an addition leads to the formation of a very low-yield stress fluid, which allows the microalgae cells to remain suspended long enough, and for the bacteria to secrete sufficient solid BC fibres, to fully encapsulate the microalgal cells (Fig. 1e(5)).

We investigated the effect of B-CNCs on the stability of microalgae solution further by adding different amounts of B-CNCs (overall B-CNCs content: 0 w/v%, 0.01 w/v%, 0.03 w/v%, 0.05 w/v%, and 0.06 w/v%) into a 4-day-old liquid culture of *C. reinhardtii* cells (OD_750_=0.5) with Tris-acetate-phosphate (TAP) medium. Fig. 2a shows that the cells alone settle to form a green pellet after just 1 day of static incubation, whilst above 0.04% w/v B-CNCs, they remain suspended in the liquid even after 30 days of static incubation.

**Fig. 2.**
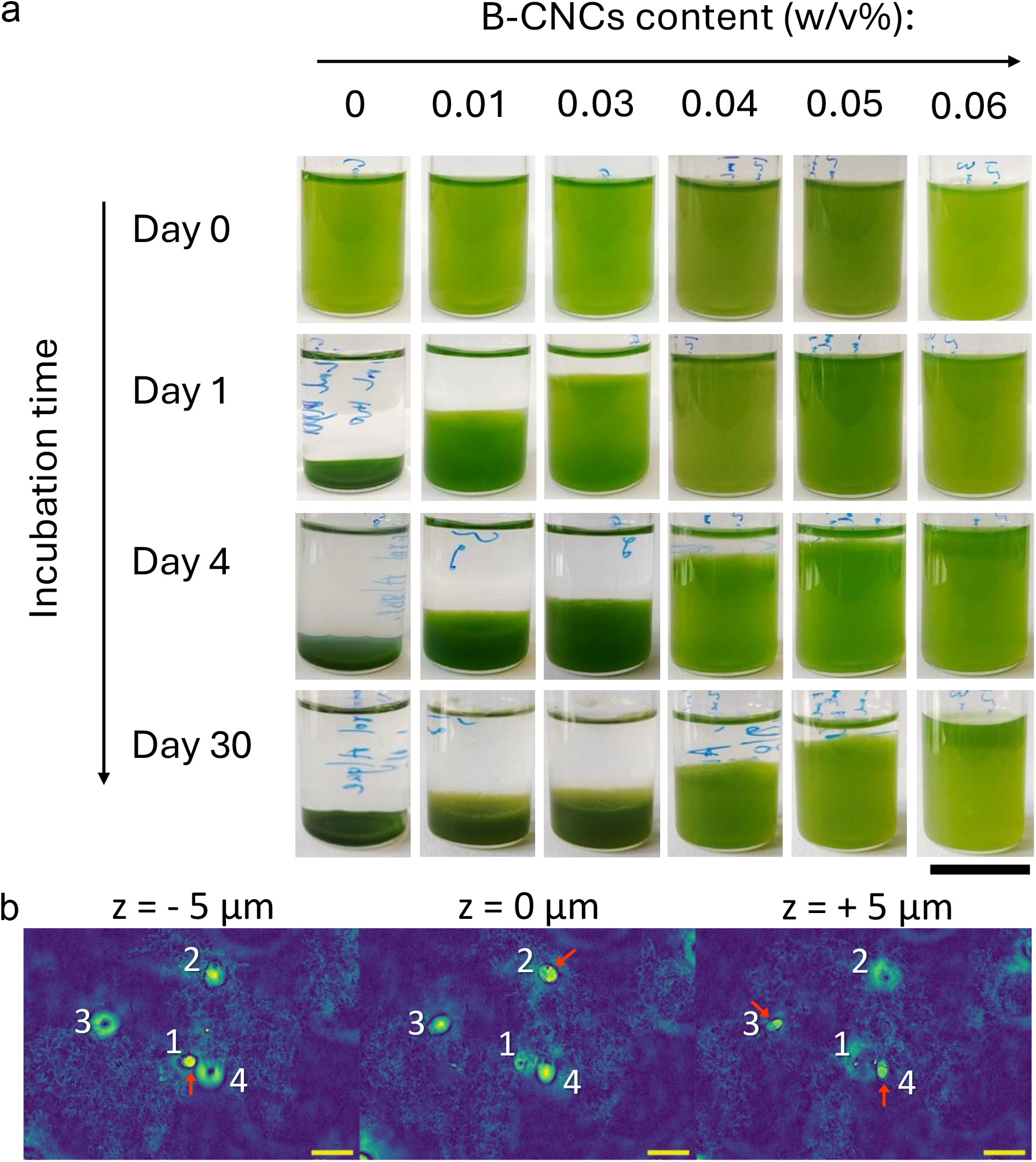
Effect of B-CNCs on the stability of microalgae solution. **a**, The microalgal suspension sediment at the bottom of the flask after 1 day of static incubation without adding B-CNCs, while the addition of B-CNCs prevented the microalgae from sedimentation (Scale bar = 2 cm). **b**, Z-stack of holotomographic images of a microalgal suspension containing 0.03% w/v B-CNCs. Red arrows indicate microalgal cells in focus in their respective frames, labelled as 1, 2, 3, and 4, and their focus positions are shown at different heights within the z-stack: cell 1 at z = -5 µm, cell 2 at z = 0 µm, and cells 3 and 4 at z = +5 µm. Clusters of CNCs are visible surrounding each microalgal cell (Scale bar = 20 µm in all images).

To study how the presence of CNCs inhibits microalgae sedimentation, we monitored the three-dimensional structure of microalgae-CNCs suspension with holotomography. Fig. 2b shows that within the microalgae-B-CNCs suspension (with 0.03 w/v% B-CNCs) microalgal cells are entrapped among loosely entangled B-CNCs clusters. Owing to this separation effect of B-CNCs on microalgal cells, the cells spread out more evenly within the suspension. This prevents the cells from clumping together or settling at the bottom, resulting in the stable suspension of microalgal cells in the liquid.

### Formulation optimization

Since microalgae cells can be stabilized within liquid in the presence of B-CNCs, a 3D bulk BC could be obtained from static culture directly with the cocultivation of bacteria and microalgae, rather than a flat thin pellicle (Fig. 3a) from the air-liquid interface with the traditional static incubation approach. To explore the optimal coculture formulation, we optimized 3 parameters (Fig. 3b-g, Supplementary Tab. 1), the B-CNCs content; the volume ratio of microalgae (TAP medium) and bacteria medium (HS medium), and the sodium hydroxide (NaOH) content.

**Fig. 3.**
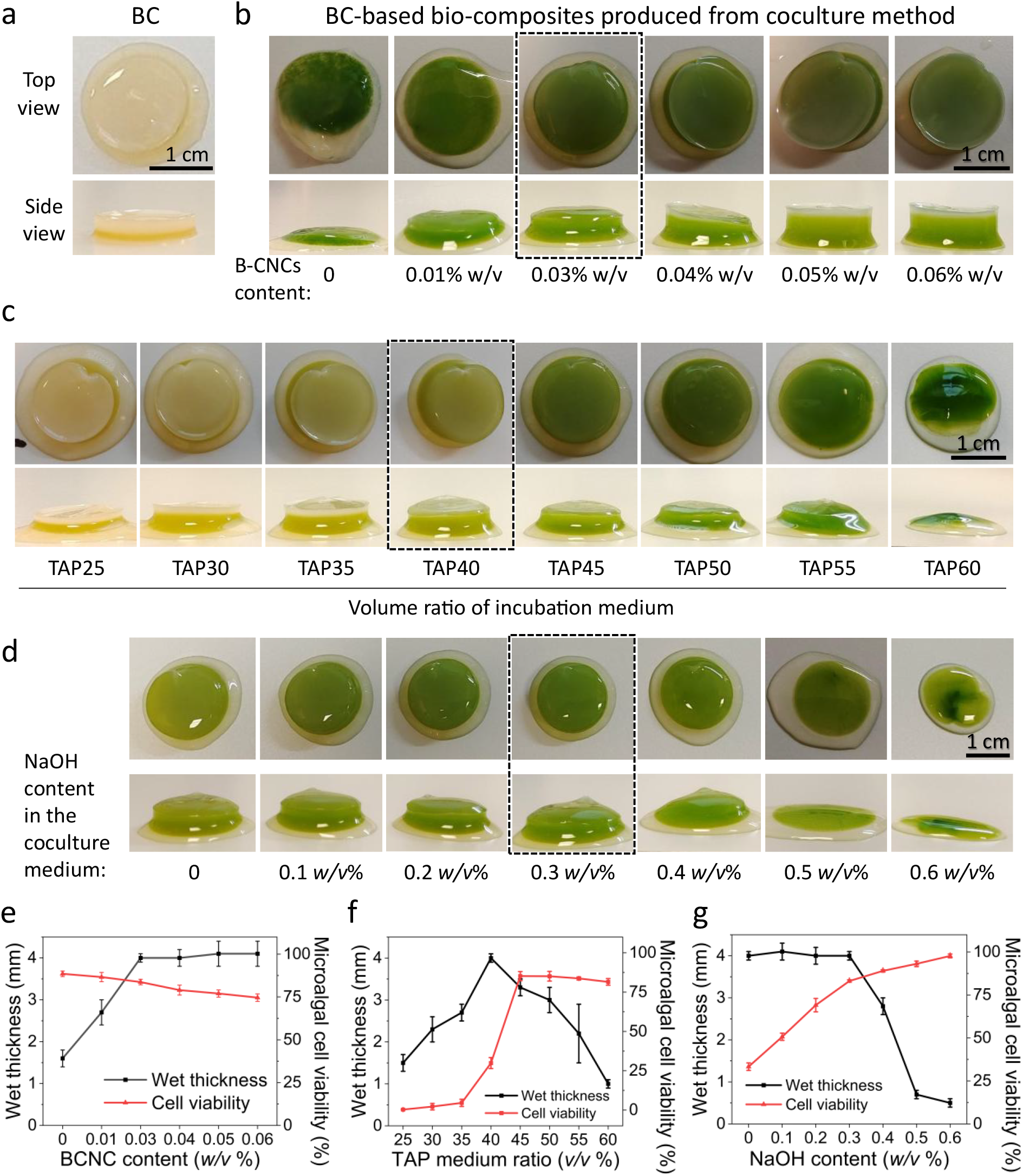
Optimisation of coculture formulation of bacteria and microalgae. **a**, Images of pure BC wet pellicle harvested from a 4-day-old *K. hansenii* static culture, where the BC pellicle formed at the air-liquid interface. **b**, The top view and side view images of BC-based bio-composites with various B-CNC content harvested from a 4-day-old *K. hansenii*/*C. reinhardtii* static coculture (with a medium volume ratio of TAP:HS=40:60, and 0.3 w/v% NaOH content). **c**, Top-view and side-view images of BC-based bio-composites with different TAP/HS medium ratios (TAP25-TAP60: The volume ratio of TAP medium) from a coculture (with a B-CNCs content of 0.03 w/v%, and no NaOH content). **d**, The top view and side view images of BC-based bio-composites with various NaOH content harvested from a 4-day-old bacteria and microalgae static coculture (with a medium volume ratio of TAP:HS=40:60 (TAP40), and 0.03 w/v% BCNCs content). The effect of **e**, B-CNC content (Supplementary Tab. 1, recipe No: 1-6), **f**, culture medium volume ratio (Supplementary Tab. 1, recipe No: 7-14) and **g**, NaOH content (Supplementary Tab. 1, recipe No: 15-21) on the wet thickness and microalgae cell viability of BC-based bio-composites.

From these results, the amount of B-CNCs clearly plays a key role in producing different wet thicknesses of BC pellicles (Fig. 3b, e). When the B-CNCs content is increased to 0.03 *w/v*% (Supplementary Table. 1, recipe No: 1-3), the wet thickness of BC-based bio-composites also increased from 1.6 ± 0.2 mm to 4.0 ± 0.1 mm, but remained unchanged when the B-CNCs content was above 0.03 *w/v*% (Supplementary Table. 1, recipe No: 4-6). This stability was attributed to the saturating effect on the stabilisation of microalgal cells with the additional inclusion of B-CNCs, and the formation of fresh BC layers without microalgae at the air-liquid interface (Fig. 3b). Further addition of B-CNCs reduced the microalgae cell viability within the BC-based bio-composites (Fig. 3e) from 83.7 ± 1.6% at 0.03 *w/v*% to 74.6 ± 2.2% at 0.06 *w/v*%. We attributed this decrease in cell viability to the increasing acidity of the culture with the presence of the negatively charged group grafted at the surface of the B-CNCs, which is unfavourable for microalgal growth^36^. Therefore, for optimal balance between microalgae cell viability and cellulose production, 0.03 *w/v* % was selected as the optimal amount of B-CNCs addition.

The second important factor influencing the wet thickness and microalgae cell viability of BC-based bio-composites (with B-CNCs content fixed to 0.03 w/v%, no NaOH added) is the bacteria (HS) and microalgae (TAP) medium volume ratio (Fig. 3c, f). When the TAP medium ratio was between 25% (TAP25) to 35% (TAP35), bacteria tended to form BC layers at the air-liquid interface (Fig. 3c), and only a thin layer of microalgae containing BC could form within the liquid, from 1.5 ± 0.2 mm (TAP25) to 2.7 ± 0.1 mm (TAP35). The microalgae cell viability remained low, from 0.3 ± 0.3% (TAP25) to 4.6 ± 2.3% (TAP35). When the TAP medium ratio was 40% (TAP40) (Supplementary Tab. 1, recipe No: 10-14), BC started to grow within the liquid, beyond the air-liquid interface. At this transition point (TAP40), the wet thickness of BC-based bio-composites reached its highest (4.0 ± 0.1 mm), and the microalgae cell viability significantly (29.9 ± 3.5%) increased when compared to TAP35 (4.6 ± 2.3%). Further increase of TAP medium ratio results in the reduction of BC production due to the limited amount of HS medium, reflected by the decrease of the wet thickness of BC-based bio-composites, from 3.3 ± 0.2 mm (TAP45) to 1.0 ± 0.1 mm (TAP60). Although the cell viability for TAP45 was higher (85.2 ± 2.9%) than that of TAP40 (29.9 ± 3.5) we decided to use TAP40 as the medium ratio to obtain higher BC production to achieve better mechanical properties of the final material.

Another parameter affecting the growth of the system was the presence of NaOH in the growing medium, (Fig. 3d, g). The formulation of co-culture nutrient media is, therefore, critical in sustaining the healthy proliferation of both species and ensuring consistent cellulose production. However, there is an inherent trade-off in creating a growth environment that accommodates both species. For example, maximal cellulose production by bacteria was observed at a low pH of around 6, while optimal photosynthesis occurs between pH 7 and 8^14^. The microalgae cell viability increased from 33.3 ± 2.2% (0 *w/v*% NaOH content) to 83.1 ± 0.7% (0.3 *w/v*% NaOH content). The addition of NaOH did not influence the wet thickness (4.0 ± 0.1 mm) of BC-based composites when NaOH content was below 0.3 *w/v*% (Supplementary Tab. 1, recipe No: 15-18). However, when the NaOH content is 0.4 *w/v*% (Supplementary Tab. 1, recipe No: 19), the wet thickness of BC-based composites decreased significantly to 2.8 ± 0.2 mm. This decrease in wet thickness was mainly due to the pH increase (Supplementary Fig. 5), which was unfavourable for BC production. In combination with the microalgae cell viability and wet thickness of BC-based composites, we decided to use 0.3 *w/v*% as the optimal NaOH amount added into the co-culture system.

In summary, by considering the trade-off between microalgae cell viability and wet thickness of BC-based composites, the optimal parameters for cocultivation of BC-based bio-composites were: 0.03 *w/v*% of B-CNC addition amount, 40% of the TAP medium ratio and 0.3 *w/v*% of NaOH content. Under this formulation, BC-based bio-composites with both high cell viability (∼ 83%) and high wet thickness (∼ 4.0 mm) were achieved.

### Daily monitoring of BC-based bio-composite growth

Fig. 4a shows the changes in the BC-based bio-composite over the first four days of development. We observed that over this time the symbiotic material grew into a green hydrogel-like composite. Microscopy indicated the *C. reinhardtii* cells and *K. hansenii* were trapped in the BC network (Fig. 4a, b) and covered by BC fibres (Fig. 4c), confirming the results from holotomography (Fig. 2b). Importantly, by analysing the liquid media surrounding the bio-composite, we observed that no cells were present by day 2, demonstrating that the cells were trapped by the BC-based bio-composites (Fig. 4a, surrounding liquid medium). This is in contrast to conventional hydrogel encapsulation^37,38^, where cell leakage into the medium remains a problem. After incubation, B-CNCs were embedded into the composite network, with no B-CNCs in the liquid media (Fig. 4a, b, c).

**Fig. 4.**
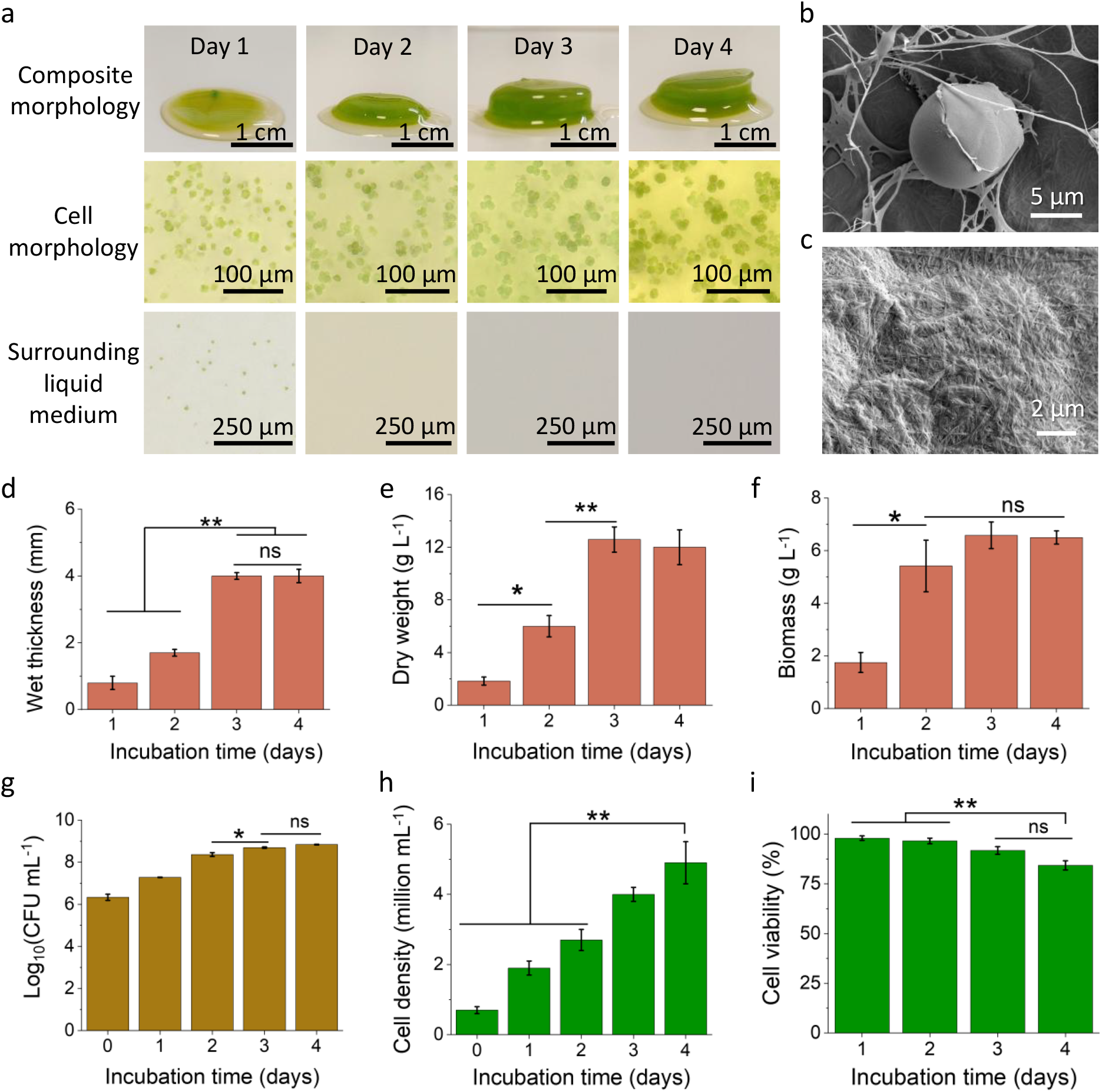
Characterization of material morphology and biological activity within BC-based bio-composites. **a**, The morphology of composite material, morphology of *C. reinhardtii* microalgae entrapped and liquid medium surrounding BC-based bio-composites in the coculture procedure, show that the material can be fabricated within 4 days of incubation, without cell leakage into the surrounding liquid medium. **b, c**, Scanning electron microscope (SEM) images of *C. reinhardtii* microalgae entangled in the fibrous network (b) and covered by BC fibers and *K. hansenii* bacteria (c). **d**-**f**, The wet thickness (d), dry weight (e) and biomass (f) values of the coculture samples during the incubation procedure. **g**, *K. hansenii* bacteria viability in the coculture procedure. **h, i**, Microalgae cell density (h) and cell viability (i) in the coculture procedure. (ns, not significant; **p < 0.01; *p < 0.05, as determined by one-way (single factor) ANOVA with post-hoc Tukey’s HSD)

To quantify the material growth rate, we monitored the wet thickness of BC-based bio-composites day by day (Fig. 4d). The wet thickness increased from 0.8 ± 0.2 mm (day 1) to 4.0 ± 0.1 mm by day 3, and thereafter did not change. We also monitored the dry weight of BC-based bio-composites (m_*dry weight*_) and the biomass weight for bacteria and microalgae within the BC-based bio-composites (m_*biomass*_).

The m_*dry weight*_ (Fig. 4e) increased in the first three days, and remained constant on day 4, similarly for m_*biomass*_ (Fig. 4f). We observed that mass of cellulose produced by bacteria ( m_*dry weight*_ − m_*biomass*_) peaked on day 3 and slowed on day 4. In the final grown BC-based bio-composites (dry state), the overall BC content was 45.8w/w% while the biomass content was around 54.2w/w%. By calculating the BC production yield, we could conclude that over 94.5% BC was produced within 3 days of incubation, with an overall BC production yield of 5.2 g L^-1^, which is about 20% higher than what has been reported for pure BC (4.3 g L^-1^) ^19^.

To understand the cell behaviour during the coculture process, we monitored the cell viability of both microalgae and bacteria. The viability of *K. hansenii* (Fig. 4g) increased from 6.3 ± 0.1 Log(CFU mL^-1^) (day 0) to 8.8 ± 0.01 Log(CFU mL^-1^) (day 4). The cell density of *C. reinhardtii* (Fig. 4h) increased from 0.7 ± 0.1 million mL^-1^ (day 0) to 4.9 ± 0.6 million mL^-1^ (day 4). The viability of *C. reinhardtii* (Fig. 4i) decreased from 98.0 ± 1.1% (day 1) to 84.3 ± 2.3% (day 4). Although the viability of *C. reinhardtii* dropped after 4 days of incubation, this value was still as high as 80 %. By monitoring the growth of coculture, we concluded that cellulose production reached the stationary phase after 4 days of incubation. Therefore, we use 4 days as the final incubation time to grow the BC-based bio-composites.

### Fed-batch culture and cell storage

To assess the long-term viability of *C. reinhardtii* encapsulated within the cellulosic matrix, we investigated the potential to perform a fed-batch culture, following a period of nutrient deficiency. Once the coculture was completed after 4 days of incubation, the bio-composites were transferred into fresh TAP medium. After 1 day of incubation in TAP medium, the BC-based bio-composites turned greener (Fig. 5a), indicating that *C. reinhardtii* maintained a constant physiological state in terms of growth rate. The *C. reinhardtii* cells formed aggregations after incubation in TAP medium (Fig. 5b). The formation of aggregations was mainly due to the movement of microalgae cells being restricted by BC fibres, resulting in limited space for the freshly divided microalgae cells to occupy. To adapt to environmental stress, these microalgae cells form aggregates, similar to what is observed when encapsulated in hydrogels^2,29,39,40^.

**Fig. 5.**
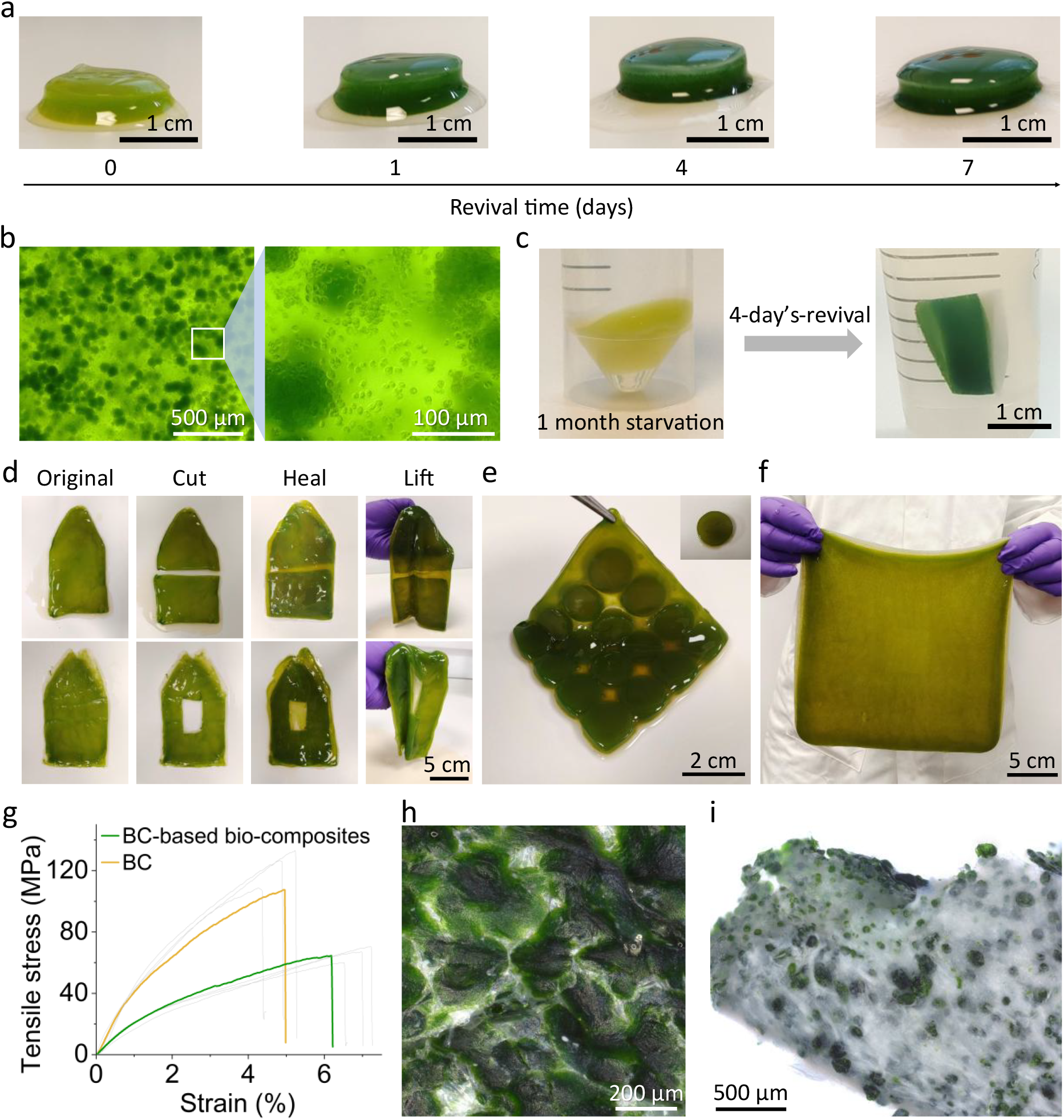
Advantages of BC-based bio-composites. **a**, BC-based bio-composites after 1, 4 and 7 days of incubation with TAP medium. The material turned greener since day 1. **b**, Optical microscopic images of BC-based bio-composites (wet) after 4 days of continued incubation in TAP medium, during which microalgal cells form aggregations. **c**, Cell storage within BC-based bio-composites. The embedded microalgae continued growing by supplementing with TAP medium after 1 month of storage without nutrient supply. **d**, The regenerative properties of BC-based bio-composites. The material could be regenerated by further incubation after being cut into two pieces or drilled a hole. **e, f**, Scalability of BC-based bio-composites. The pre-grown building blocks of BC-based bio-composites could be further incubated into a larger piece of material (e). Larger BC-based bio-composites can be directly grown from the direct incubation of coculture solution in a larger vessel (f). **g**, Comparison of tensile properties, showing similar mechanical properties between dried BC film and dried BC-based bio-composite film. **h**, Surface morphology of dried BC-based bio-composites, showing a rough surface. **i**, Microalgae cell morphology in dried BC-based bio-composites, forming cell aggregations within the sample.

To assess the potential of the BC-based bio-composites in preserving cell viability during storage (Fig. 5c), the 4-day-old cocultured material was placed in an empty falcon tube, without additional medium except for the moisture retained by the BC-bio-composite. The falcon tube was sealed and kept at ambient laboratory conditions for 1 month, and then 30 mL TAP medium was added to check the status of the algal cells. Within 4 days, the sample became green, indicating the promising potential of the BC-based bio-composites in preserving cell viability during storage without nutrient supplements. The long-term viability of cells is mainly due to the hydrophilic nature of BC networks. This feature could facilitate shipping, or space colonisation without being damaged^29^.

### Regeneration and scalability

Apart from the potential for cell storage, the BC-based bio-composite material produced has self-healing characteristics: Fig. 5d shows that the bio-composites can regenerate when adding a fresh coculture liquid mixture (TAP-HS) after the composite is cut in two, allowing the formation of fresh material between the junction points, rejoining the two pieces. Such properties can also be exploited to assemble pre-grown building blocks into more complicated 3D shapes with further incubation (Fig. 5e).^9^ Furthermore, by growing the material directly in larger vessels, it is possible to produce a scalable and homogeneous film on a larger scale (Fig. 5f).

### Mechanical stability

To compare the mechanical properties of BC-based bio-composites with pure BC, tensile tests were performed. As the water content influences the accuracy of tensile test results, we decided to perform the tensile tests under dry conditions for a fair comparison of the mechanical properties. The tensile test results (Fig. 5g) showed that after drying, BC-based bio-composites had a higher elongation at break value (6.7 ± 0.2%) and a lower tensile strength value (65.5 ± 2.2 MPa) compared to that of a pure BC film (elongation at break: 4.8 ± 0.2%; tensile strength: 118.9 ± 6.4 MPa). The increase in elongation at break was due to the special structure of dried BC-based bio-composites film, where a rough sample surface (Fig. 5h) was formed during the drying process, making the sample extend longer under tension.^23^ The reduction in tensile strength is probably attributed to the presence of microalgae cells, which formed aggregations during incubation (Fig. 5i). Due to the presence of these microalgal cells, the BC fibre density (Supplementary Fig. 7) within the same cross-section area was smaller than pure BC, making BC-based bio-composites less strong.^41^ Nevertheless, BC-based bio-composites still demonstrated mechanical robustness comparable to pure BC. For instance, a filament of the bio-composite with a diameter of 2 mm was able to lift a weight of 2 kg (Supplementary Fig. 8).

## Conclusion

In summary, we have demonstrated the production of bulk BC, and enabled the facile fabrication of BC-based bio-composites with complex 3D geometries, through simple static incubation of the BC-producing bacteria with the microalga *C. reinhardtii*, eliminating the need for special templates or complicated processing techniques^10,12,26,27^.

We observed that the presence of CNCs, (derived from both BC as well as plant sources), allows the creation of a very low-yield stress fluid that avoids the precipitation of the microalgae (oxygen-generating sites) and bacteria within the liquid medium^42,43^.

By optimising the formulation (0.03 *w/v*% of B-CNCs addition amount, 40% of the TAP medium ratio and 0.3 *w/v*% of NaOH content), we obtained BC-based bio-composites with both high cell viability (∼ 83%) and excellent mechanical robustness (comparable to BC), which is challenging to achieve in engineered living bulk hydrogels^29,32,33^. Moreover, the BC-based bio-composites can trap all the microalgal cells from the medium into the freshly formed BC network without cell leakage only after 2 days of incubation, which could be possibly expanded in the context of wastewater treatment^38,44- 46^.

In the composite, the overall dry content for BC was 45.8w/w% and the biomass was around 54.2w/w%, so we believe that such systems could also be optimised for use as next-generation photobioreactors^47^, reducing the energy cost and environmental pressure from the traditional downstream processing of other microalgal liquid cultures^30,36^. For BC fibres within the BC-based bio-composites, over 94.5% BC was produced freshly within 4 days of incubation, with an overall BC production yield of 5.2 g L^-1^, which is higher than pure BC (4.3 g L^-1^) in monoculture of *K. hansenii*^19^.

Overall, this BC-based bio-composite possesses many advantages, including tunable and flexible geometry, high cell viability, excellent mechanical robustness, low cell leakage, cell storage ability, regenerative properties and scalability, enabling its broad applications in engineered living materials^32^, photobioreactors^30,47^ and the tissue engineering field.^48^ Apart from these applications, this coculture method opens a new window in fabricating fibre-reinforced composites by the addition of various components, such as graphene oxide, indium tin oxide, bio-glasses, etc., into the coculture solution through direct static microbial cultivation, providing a universal approach for the biological production of sustainable fibre-reinforced biomaterials.

## Data availability

All produced data that support the main figures of this study are included in this published article. Data points for the mechanical tests are provided as Source data files. Additional data are available from the corresponding author upon request.

## Acknowledgements

We thank R. M. Parker, R. Li, J. Wang, Q. Shen, M.-E. Aubin-Tam, for advice and discussions, we thank L. J. Occhipinti, M. Kaya, S. Wojno, L. Archer, H. Greer, for their help with some experiments. K. Y. and S. V. were funded by ERC BiTe ERC -2020 -CoS-101001637. S. T. C. was funded by the Harding Distinguished Postgraduate Programme. SV and AS acknowledge the SNSF Sinergia 198750 grant. The TEM operation was supported by EPSRC Underpinning Multi-User Equipment Call (EP/P030467/1).

## Competing interests

The authors declare no conflict of interest.

## Methods

### Strains and chemicals

Cellulose producing bacterium *Komagataeibacter hansenii* ATCC 53582 (*K. hansenii*) was provided by Prof. T. Ellis, microalga *C. reinhardtii* WT12 (CC124) was obtained from the Chlamydomonas Resource Center at the University of Minnesota (https://www.chlamycollection.org/). All chemicals, unless otherwise specified, were purchased from Sigma-Aldrich.

### Bacterial cellulose (BC) incubation and purification

BC solid pellicle was prepared by inoculating 1 *v/v*% *K. hansenii* into Hestrin–Schramm (HS) medium (tryptone: 5.0 g L^−1^, yeast extract: 5.0 g L^−1^, disodium hydrogen phosphate: 2.7 g L^−1^, citric acid: 1.5 g L^−1^, and glucose: 20 g L^−1^) under static incubation at 30 °C for 4 days. A solid BC pellicle harvested at the air/liquid interface was purified by boiling within 1 w/v% sodium hydroxide (NaOH) solution for 20 mins for 3 times, followed by rinsing with deionised water to remove the unreacted medium within BC until the solid pellicle turned white.

### Preparation of bacterial cellulose nanocrystals (B-CNCs)

To prepare B-CNCs, we first freeze-dried the purified wet BC pellicles ( BenchTop Pro SP Scientific freeze-drier) at -76.6 °C under a low pressure of 23 mT, and ground them into flakes with a Cookworks grinder (Multiblender mill, Nihonseiki Kaisha, ltd, BLAS-501, 220-240 V∼50 Hz 150 W) for 1 min. This was followed by adding the BC flakes into 60 *wt*% sulfuric acid with an acid/cellulose ratio of 70 mL g^-1^, under constant stirrig at 500 rpm and 60 °C. B-CNCs formed after this BC hydrolysis procedure which lasted for 1 h.

After acid hydrolysis, the B-CNCs was poured into 3 times volume of Milli-Q water to quench the reaction. The B-CNCs suspension was then centrifuged with a BECKMAN COULTER (Avanti J-26 XPI) centrifuge (JLA-81000 rotor, 15900 g, 4 °C, 15 mins) for 3 times to remove the acid. After centrifugation, the solid B-CNCs was resuspended into 100 mL Milli-Q water. To further remove the acid residuals, the B-CNCs solution was transferred into a dialysis tube (Medicell Membranes ltd, Visking code DTV12000.11.15, 38.1 mm, 12-14000 Daltons), and the water was changed daily for a week.

To increase the stability of B-CNCs dispersion, tip sonication was carried out under ice bath, Amplitude=30%, 2s on, 1s off, with an effective sonication time of 5 mins, and a total sonication time of 7 min 30 s. The sonicated B-CNCs solution was then filtered with an 8-µm pore size filter paper to remove the large aggregates. The B-CNCs solution was then autoclaved and stored in a Simax bottle at 4 °C for use.

### Morphological characterisation

The morphology of B-CNCs was examined using scanning electron microscope (SEM) and transmission electron microscope (TEM). To prepare the SEM sample, the B-CNCs suspension was first diluted by a factor of 10^6^. Then, 50μL of the diluted B CNCs suspension was drop-casted onto carbon tape and allowed to dry at 60 °C for two days. The samples were then sputter-coated to a nominal thickness of 10 nm with platinum (Quorum Q150T ES). SEM was performed using an acceleration voltage of 5 kV and a working distance of 3-6 mm under a TESCAN MIRA3 FEG-SEM system. For TEM, 10 μL of diluted B-CNCs suspension was drop-casted onto a carbon-coated copper grid (Agar Scientific S160-3 Carbon film 300 mesh Cu) prepared by glow discharge and stained with uranyl acetate solution. TEM was operated using a Talos F200X G2 microscope (Thermo Scientific, FEI) at accelerating voltage of 200 kV and a CCD camera.

### Preparation of microalgae liquid culture

*C. reinhardtii* cells were maintained on agar slopes in Tris-acetate-phosphate (TAP) medium^49^. To prepare liquid cultures, we inoculated 1 *v/v*% *C. reinhardtii* (OD_750_=0.5) into TAP medium and incubated for 4 days in a Panasonic MLR-352-PE incubator at 25 °C with LED light (800 W) under 18h/6h light/dark conditions.

### Incubation of BC-based bio-composites

To prepare the algal-bacterial coculture, a 4-day-old *C. reinhardtii* liquid culture (50 mL) was centrifuged (Hettich ROTINA 38R) at 1780 g for 5 mins at 20 °C, then the microalgae solid pellet was resuspended into fresh TAP medium with an optical density (OD) absorbance value of 0.5 at 750 nm, with fresh TAP medium as reference, measured with an JENWAY 6300 Spectrophotometer. The supernatant of 4-day-old bacteria culture, microalgae liquid culture (OD_750_=0.5, exponential phase), B-CNCs, and NaOH solution were mixed together with an overall composition of 0.03 *w/v*% B-CNCs, 40 v*/v*% of the TAP medium ratio and 0.3 *w/v*% of NaOH content. For other recipes to explore the coculture parameters, the respective amount was added accordingly (Table 1). All samples for formulation optimisation were incubated in 12-well plates (no coatings) with an overall volume of 4 mL coculture solution. All samples were incubated in static in a Panasonic MLR-352-PE incubator at 25 °C with 3Ls light (800 W) under 18h/6h light/dark conditions for 4 days.

### Determination of cell density and viability

The morphology of BC-based bio-composites was monitored with a Keyence VHX-7000 digital optical microscope in transmission mode. The wet thickness of BC-based bio-composites was measured with a calliper.

To measure the microalgae cell density and cell viability within BC-based bio-composites, we first transferred the BC-based bio-composites from a 12-well plate to a 50 mL falcon tube, followed by the addition of μL cellulase from *Trichoderma reesei* (aqueous solution, ≥700 units g^−1^). The falcon tubes were then transferred to a INFORS HT shaking incubator at 30 °C, 250 rpm for 4 hours. After the cellulase treatment, BC-based bio-composites were dissolved and the cells were suspended freely in liquid medium. We then centrifuged the cell suspension (Hettich ROTINA 38R) at 1780 g for 5 mins at 20 °C to remove the cellulase and liquid medium. The solid cell pellet was then resuspended into 4 mL TAP medium for subsequent characterisation.

To measure the microalgal cell density, we added 10 μL of this resuspended liquid into a, Marienfeld Neubauer hemocytometer (0.100 mm in depth, 0.0025 mm^2^ area) and determined the cell numbers under a Keyence VHX-7000 digital optical microscope under a 400× magnification. To measure the microalgal cell viability, we mixed this resuspended liquid and a 0.4% Trypan Blue solution (Gibco) with a volumetric ratio of 1:1 for 5 mins. The mixture was centrifuged with a JENCONS -PLS 24D Spectrafuge for 5 mins at 13000 rpm, 20 °C. The supernatant containing the Trypan Blue dye was removed and same amount of TAP medium was added to resuspend the microalgal cells. The microalgal cell viability was determined with a hemocytometer as before; dead microalgal cells appeared blue, while the living microalgae cells were green. The proportion of viable cells was determined in biological and technical triplicates.

To measure the bacterial viability within BC-based bio-composites, we took the cocultured BC-based bio-composites with different incubation time from 12-well plate into 50 mL falcon tube and repeated the above-mentioned cellulase treatment. Afterwards, the *K. hansenii* bacterial viability was assessed with the colony-forming unit (CFU) measurements. Briefly, the bacterial pellet was resuspended in the same initial volume of saline (0.9% w/v NaCl). We then prepared dilutions ranging from 10^0^ to 10^−8^, followed by spotting 10 μL of each dilution onto HS-agar plates (supplemented with 2% *v/v* acetic acid) in biological triplicates. The HS-agar plates were then incubated at 30 °C for 3 days, and the number of colonies was enumerated and the log_10_ (CFU mL^-1^) was calculated in biological and technical triplicates^19^.

### Tomocube holotomography

Three-dimensional holotomograms of individual cells were reconstructed using low-coherence illumination from various angles. The Holotomography system (HT-X1, Tomocube Inc., Daejeon, Republic of Korea) employs an LED with a central wavelength of 450 nm for stable illumination. This system includes a specially designed condenser lens with a numerical aperture (NA) of 0.72 and a long working distance of 30 mm. It is paired with a motorised module and a high numerical aperture objective lens (NA 0.95, UPLXAPO40X, Olympus) to capture multiple intensity stacks from thick specimens under different modulated illuminations. These stacks are then processed to reconstruct the 3D holotomogram of the sample.

### Dry weight determination

To measure the total dry weight of BC-based bio-composites (m_*dry weight*_), we transferred the 4-day-old BC-based bio-composites from 12-well plates into falcon tubes, followed by drying in an oven at 40 °C for 4 days. To measure the dry weight of biomass within the BC-based bio-composites (m_*biomass*_), the BC-based bio-composites were subjected to the same treatment with cellulase as aforementioned to remove the cellulosic matrix. The resulting cell pellets (bacteria and microalgae) were washed and dried in an oven at 40 °C for 4 days. The BC yield within the BC-based bio-composites can be calculated with the following equation:

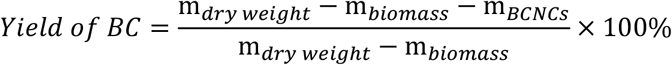

where m_*dry weight*_ is the dry weight of BC-based bio-composites, including the BCNCs, microalgae, bacteria and freshly formed BC; m_*biomass*_ is the weight for bacteria and microalgae; m_*BCNCs*_ is the initial amount of BCNCs added to the coculture solution.

### Revivability and longevity of BC-based bio-composites

To determine if the microalgal cells embedded within the BC-based bio-composites can be revived, we transferred the 4-day-old BC-based bio-composites into a 50 mL falcon tube with 30 mL TAP medium. Then we incubated these falcon tubes in a Panasonic MLR-352-PE incubator at 25 °C with 3Ls light (800 W) under 18h/6h light/dark conditions for 1, 4 and 7 days. To check the longevity of BC-based bio-composites, we first transferred the 4-day-old BC-based bio-composites into an empty 50 mL falcon tube, then we closed the lid and kept the falcon tube at ambient lab conditions for 1 month. Afterwards, we added 30 mL TAP medium into the falcon tube for revival.^29^

### Regenerative properties of BC-based bio-composites

To check the regenerative properties of BC-based bio-composites, we first prepared the BC-based bio-composites by adding 100 mL coculture solution into a 50 ml cell culture flask, then we incubated the flasks in a Panasonic MLR-352-PE incubator at 25 °C with LED light (800 W) under 18h/6h light/dark conditions for 4 days. Afterwards, we took the BC-based bio-composites out of the incubation flask and created damage by cutting the samples into 2 pieces or digging the sample with a hole. We transferred these damaged samples into a freshly sterilized cell culture flask and healed the samples by adding 50 mL coculture solution into a 50 ml cell culture flask. Then flasks were incubated again in static in a Panasonic MLR-352-PE incubator at 25 °C with LED light (800 W) under 18h/6h light/dark conditions for 4 days. The damaged BC-based bio-composites can be healed after incubation.

### Scalability of BC-based bio-composites

To prepare scalable BC-based bio-composites, we used two approaches: In the first approach, we directly pour 300 mL coculture solution into a 30*30 cm glass vessel, and then we incubated the vessel in static in a Panasonic MLR-352-PE incubator at 25 °C with LED light (800 W) under 18h/6h light/dark conditions for 4 days. In the second approach, we first prepared many building blocks by growing 4 mL coculture solution in 12-well plates in static in a Panasonic MLR-352-PE incubator at 25 °C with LED light (800 W) under 18h/6h light/dark conditions for 4 days, followed by further growing these building blocks into a larger piece into a 50 ml cell culture flask with the addition of 50 mL coculture solution under same conditions.

### Tensile properties of BC-based bio-composites

To check the tensile properties of BC and BC-based bio-composites, we added 100 mL bacterial monoculture or coculture solution into a 50 ml cell culture flask, then we incubated these flasks in static in a Panasonic MLR-352-PE incubator at 25 °C with LED light (800 W) under 18h/6h light/dark conditions for 4 days. Afterwards, we took these samples out of the flasks, put them overtop of the flasks and dried them in Class II cabinet for 5 days. After drying, we cut them into a dimension of 60 mm* 10 mm*0.05 mm (Pure BC), 60 mm* 10 mm*0.1 mm (BC-based bio-composites) for tensile testing. The tensile tests were performed using an Instron universal testing machine with a 10 kN load cell and 1 kN grips. The measuring distance between the clamps was 20 mm, and the samples were tested with a loading rate of 5 mm min^-1^. At least three specimens per group were measured for the data presented in this work.

### Statistics

Statistical analysis was performed on https://astatsa.com/. The experimental groups were compared using one-way (single factor) ANOVA with post-hoc Tukey's HSD (honest significant difference) tests^19^.

## Notes

### Competing Interest Statement

The authors have declared no competing interest.

