## Supplementary information for "Cultivating Future Materials: Artificial Symbiosis for Bulk Production of Bacterial Cellulose Composites"


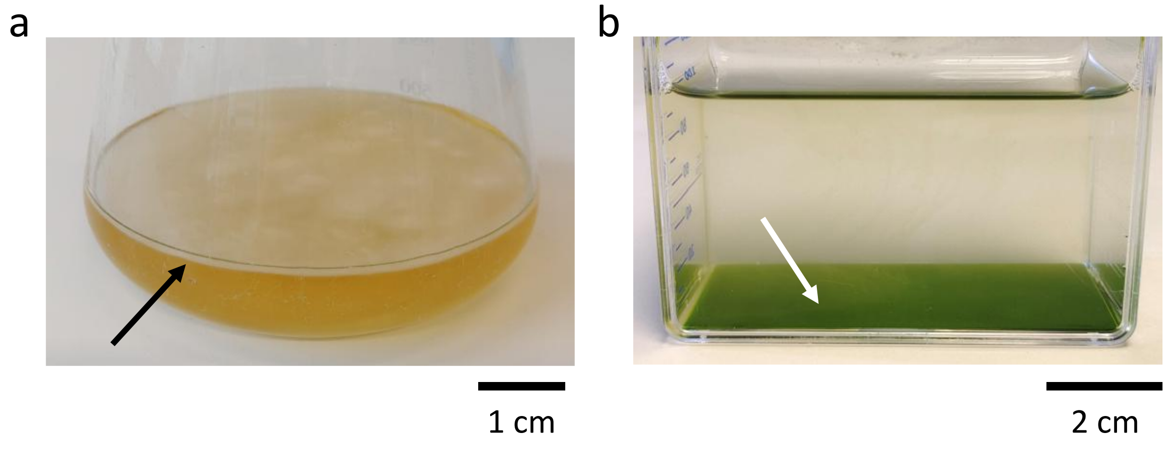


**Supplementary Fig. 1 | Photographs of bacteria (*K. hansenii*) and microalgae (*C. reinhardtii*) liquid culture after 4 days of static incubation. a**, A solid bacterial cellulose (BC) pellicle (black arrow) forms at the air-liquid interface. **b**, Microalgae cells (white arrow) sediment at the bottom of the incubation flask.

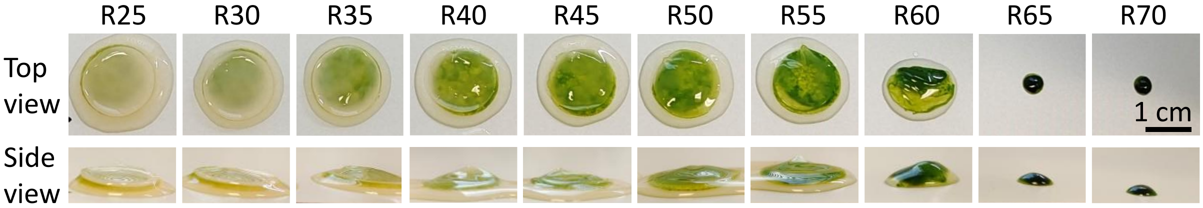


**Supplementary Fig. 2 | Photographs of BC-based bio-composites through the cocultivation of bacteria (*K. hansenii*) and microalgae (*C. reinhardtii*).** *K. hansenii* bacteria and *C. reinhardtii* microalgae liquid culture with different volume ratio (e.g. R25: The medium ratio of microalgae: bacteria =25:75) are incubated under static for 4 days. In most cases, only a thin layer of microalgae-BC bio-composites (green part) forms and attaches at the bottom side of BC, as the result of the sedimentation of microalgae cells during the coculture procedure.

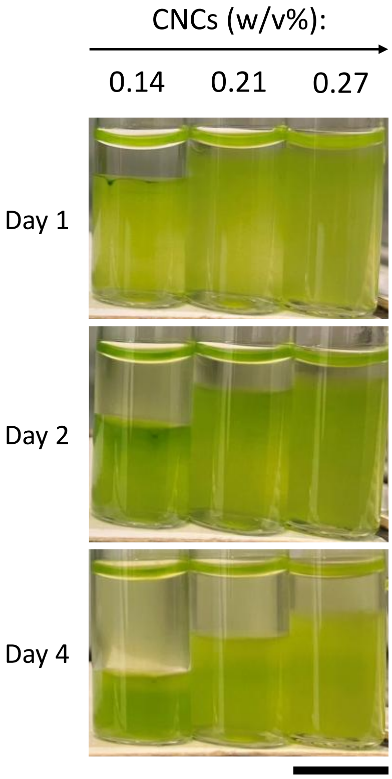


**Supplementary Fig. 3 | Effect of CNCs on the stability of microalgae solution.** The microalgal suspension sediment more slowly with the addition of CNCs (Scale bar = 1 cm).

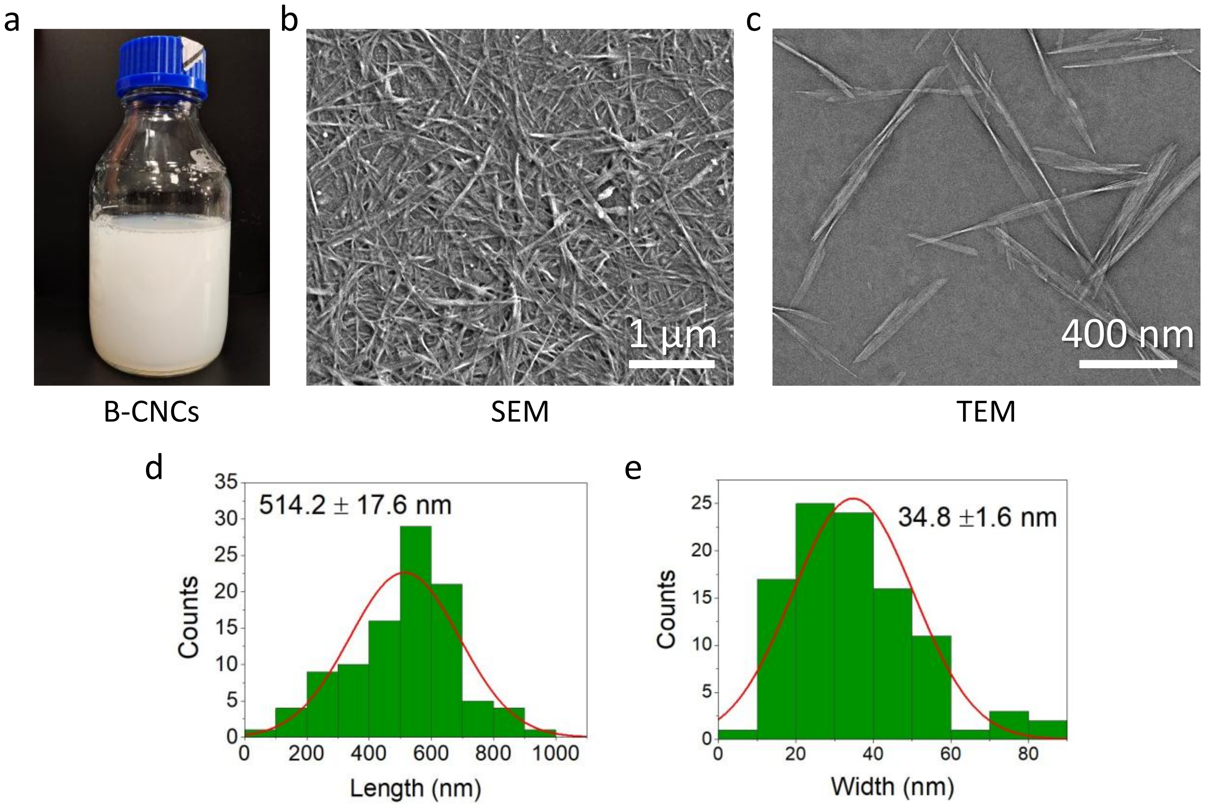


**Supplementary Fig. 4 | Morphology of bacterial cellulose nanocrystals (B-CNCs). a**, Photograph of B-CNCs suspension, which remains suspended and stable even after several months. **b**, SEM and **c**, TEM image of B-CNCs, showing a needle-like morphology. **d**, Length and **e**, width distribution of B-CNCs based on TEM images.

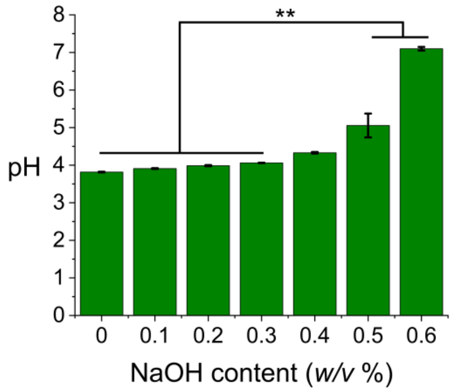


**Supplementary Fig. 5 | pH change of microalgae and bacteria coculture solution with different NaOH content.** When the addition of NaOH is above 0.4 w/v%, pH started to increase significantly (* $p< 0.05$; ** $p<0.01$ as determined by one-way (single factor) ANOVA with post-hoc Tukey's HSD).

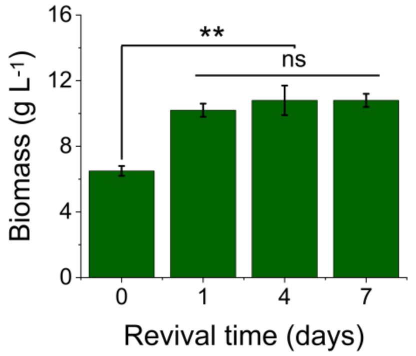


**Supplementary Fig. 6 | Change of biomass values for BC-based bio-composites after being revived in TAP medium.** The biomass of BC-based bio-composites increased from 6.5 ± 0.3 g $L^{-1}$ (Day 0) to 10.2 ± 0.4 g $L^{-1}$ (Day 1) and remained unchanged in day 4 (10.8 ± 0.9 g $L^{-1}$) and day 7 (10.8 ± 0.4 g $L^{-1}$), representing that 1 day is sufficient for the revival process.

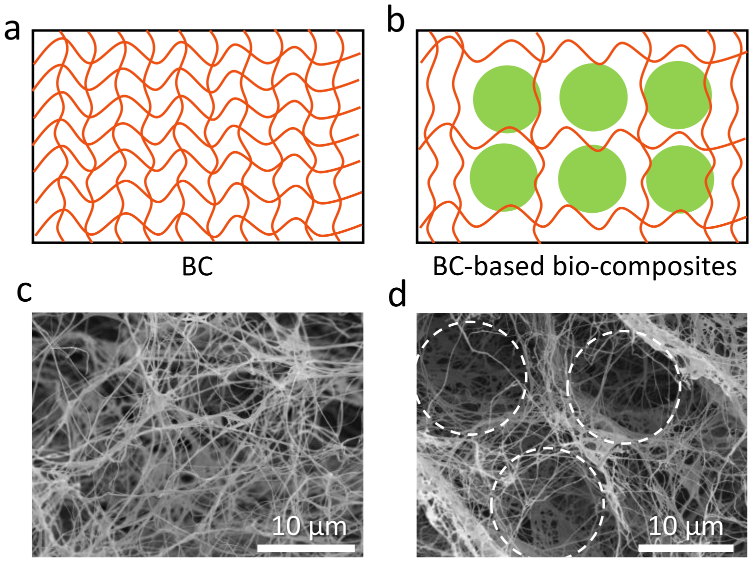


**Supplementary Fig. 7 | Comparison of fiber density between BC and BC-based bio-composites.** Illustration of fiber density for **a**, BC and **b**, BC-based bio-composites. SEM images of **c**, BC and **d**, BC-based bio-composites after drying, the white circles were spaces occupied by microalgae cells, resulting in the reduced fiber density.

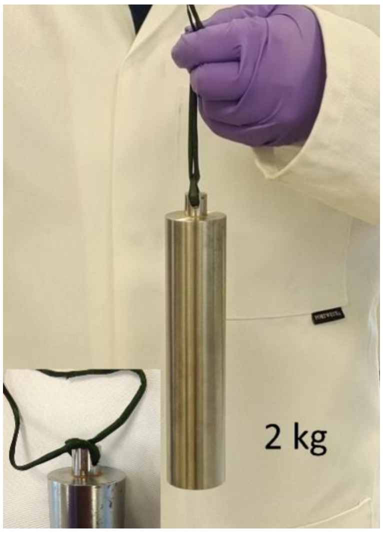


**Supplementary Fig. 8 | Mechanical robustness of BC-based bio-composites.** A 2 kg weight can be lifted with the BC-based bio-composites filament, representing its high robustness.

**Supplementary Tab. 1** Recipe composition during formulation optimization

| Recipe No. | Recipe Composition | | |
|---|---|---|---|
| | B-CNC content (w/v%) | Volume ratio of TAP:HS | NaOH content (w/v%) |
| 1 | 0 | 40 : 60 | 0.3 |

| | | | |
|---|---|---|---|
| 2 | 0.01 | 40 : 60 | 0.3 |
| 3 | 0.03 | 40 : 60 | 0.3 |
| 4 | 0.04 | 40 : 60 | 0.3 |
| 5 | 0.05 | 40 : 60 | 0.3 |
| 6 | 0.06 | 40 : 60 | 0.3 |
| 7 | 0.03 | 25 : 75 | 0 |
| 8 | 0.03 | 30 : 70 | 0 |
| 9 | 0.03 | 35 : 65 | 0 |
| 10 | 0.03 | 40 : 60 | 0 |
| 11 | 0.03 | 45 : 55 | 0 |
| 12 | 0.03 | 50 : 50 | 0 |
| 13 | 0.03 | 55 : 45 | 0 |
| 14 | 0.03 | 60 : 40 | 0 |
| 15 | 0.03 | 40 : 60 | 0 |
| 16 | 0.03 | 40 : 60 | 0.1 |
| 17 | 0.03 | 40 : 60 | 0.2 |
| 18 | 0.03 | 40 : 60 | 0.3 |
| 19 | 0.03 | 40 : 60 | 0.4 |
| 20 | 0.03 | 40 : 60 | 0.5 |
| 21 | 0.03 | 40 : 60 | 0.6 |
